# Anti-Inflammatory Potential of Selected Indonesian Phytogenic Blends to Nitric Oxide Synthase – Inducible Protein in Mojosari Ducks (*Anas javanica*): In-silico Study

**DOI:** 10.1101/2023.04.26.538405

**Authors:** Muhammad Andika Yudha Harahap, Cindy Audina Damayanti, Syahputra Wibowo, M. Halim Natsir, Osfar Sjofjan

## Abstract

Inflammation is a defensive response to tissue damage, infectious agents, and injury. Necrotic enteritis is an inflammatory response induced by pathogenic bacteria invading the intestines of Mojosari ducks (*Anas javanica*). In contrast, excessive nitric oxide production by inducible nitric oxide synthase during inflammatory processes can cause significant intestinal tissue damage and cellular toxicity. Oxyresveratrol is an active compound of *Morus alba* that has been known to have antioxidant activity and can suppress the inflammatory process, inhibiting the increased expression of nitric oxide synthase (iNOS). However, unfortunately, this plant is not endemic to Indonesia, so using native Indonesian spices that can be a substitute for oxyresveratrol is necessary. The docking results from nine Indonesian phytogenic blends interacting with NOS revealed that cynaroside from the *Piper betle* L. plant might be utilized instead of oxyresveratrol as an anti-inflammatory drug via the inhibitory pathway of nitric oxide synthase protein. The docking results showed that from the nine compounds tested, it can be concluded that three compounds were found that are better than the control compound (oxyresveratrol) in terms of binding affinity (energy) and the type of hydrogen bonds bond in amino acid proteins that are equal to the amount more than all compounds tested. The three compounds are cynaroside compounds from *Piper betle* L. with a binding energy of -9.4 kcal / mol and a Conventional Hydrogen bond type GLU(B):761, GLU(A):761, curcumin compounds from *Curcuma longa* L. with a binding energy of -8.6 kcal/mol and Conventional Hydrogen bond type GLN(A):760, GLN(B):760 and compound 14-deoxy-11, 12-didehydroandrographolide from *Andrographis paniculata* with binding energy -8.8 kcal/mol and Conventional Hydrogen bond type GLU(B):761 can be used instead of oxyresveratrol as an anti-inflammatory agent through the inhibition pathway of Nitric Oxide Synthase protein (NOS)

## INTRODUCTION

Inflammation is an essential defensive response to injury, tissue damage, or infectious factors. A slow inflammatory response will lead to further tissue damage and chronic diseases such as colitis. (Majumder et al., 2016; Franceschi et al., 2014). Inflammation can be treated by regulating several macrophages, phagocytosis, and pro-inflammatory cytokines (NO, IL-6, TNF-α, IL-1β) (Laveti et al., 2013). Nitric Oxide (NO) is an important mediator of inflammation. NO is formed endogenously by the NO synthase (NOS) enzyme type by utilizing L-arginine as a substrate and oxygen molecule and NADPH as a cofactor. Most NO production is catalyzed by inducible nitric oxide synthase (iNOS). Inflammation occurs when NO and iNOS levels rise in tissues (Lukiati et al., 2012). Enteritis is one of the features of gastrointestinal diseases caused by microorganisms. Enteritis is characterized by inflammation in the intestines that can develop into Necrotic Enteritis (NE), which can aggravate inflammatory conditions in the intestines. Various avian organisms, including the Mojosari duck, can experience this disease. The Mojosari laying duck (*Anas javanica*) is a type of Indonesian laying duck with a high egg-producing ability and a role as a reservoir in the spread of the AI virus with a rapid and widespread spread due to post-harvest grazing maintenance patterns. The role of ducks as waterfowl in the natural host of AI viruses results in higher seroprevalence outcomes. Clinical symptoms accompany it, and the virus continues to be excreted for a long time, giving rise to various inflammatory diseases, including NE (Ungsyani et al., 2021). Such lesions impact the performance of ducks and cause economic losses due to a decrease in the productivity of laying ducks. Symptoms and signs of hemorrhagic conditions in the intestinal mucosa in the digestive tract of duck cause inflammation in the inner lining of the intestine, triggering the condition of hemorrhagic necrosis lamina propria to cause abscesses. An already severe acute inflammatory condition is characterized by bleeding along the gastrointestinal mucosa of laying ducks, so inhibition is needed to be capable of being an anti-inflammatory agent in the digestive tract of the duck’s intestines.

The commonly used anti-inflammatory agent is antibiotics, but the ban on antibiotics as feed additives has encouraged many studies to look for other alternatives to substitute antibiotics. One of those is phytogenic blends such as Natural Growth Promoters (NGP) which has been widely used as an effective alternative to natural antibiotics. Phytogenic blends such as NGPs are highly developed as feed additives, immunity, performance enhancing, and highly effective in improving gastrointestinal health (Panda et al., 2009). The active substance derived from herbal plants is generally found in secondary metabolites. Indonesia is a country with abundant biodiversity, including various kinds of medicinal plants. Using some medicinal plants as phytogenic blends is very common in Java Island, an area where Mojosari duck is widely farmed. Some herbal plants that are easy to find on Java Island and can be used as phytogenic blends and anti-inflammatory agents are *Curcuma longa* L., *Andrographis paniculata, Kaempferia galanga, Zingiber officinale, Phyllanthus urinaria, Piper betle* L. and *Tinospora cordifolia* which have advantages as antioxidant compounds and anti-inflammatory agents to improve the work of the digestive organs in laying ducks.

While the plant *Eleutherine palmifolia* (L.) Merr. is one of the medicinal plants from the island of Kalimantan which is also in Indonesia. The plant contains oxyresveratrol as an active compound. This compound has antioxidant activity and can suppress the inflammatory process, inhibiting the increased expression of Nitric Oxide Synthase (iNOS) stimulated by LPS. The results showed that the anti-inflammatory properties of oxyresveratrol correlated with the inhibition of iNOS expression through down-regulation of NF-μB-binding activity and significant inhibition of COX-2 activity (Iqbal et al., 2012). The oxyresveratrol compound from Dayak onions will be used as a control to compare its inhibition ability against iNOS proteins with phytogenic blends originating from Java Island. The method in this study uses protein modeling and molecular docking to demonstrate the ability to produce accurate predictions of new knowledge about how drugs work while helping to reduce the cost of developing drug candidates through in vivo testing. In-silico tests, computer software developed to model pharmacological or physiological processes is used before stepping into controlled in vitro and in vivo experiments (Chadalawa et al., 2018). The results of ligand-receptor interactions from the computational analysis will then be used as reference active compounds in this study. The comprehensive protein modeling and molecular docking method are expected to be able to explore the potential of phytogenic blends of active compounds as anti-inflammatory herbal ingredients for cases of enteritis in Mojosari laying ducks (*Anas javanica*).

## Material and Methods

### Data mining

Data mining in in-silico research was carried out by obtaining control ligands and phytogenic blends data from the PubChem database https://pubchem.ncbi.nlm.nih.gov/ (PubChem ID for control, namely oxyresveratrol (5281717), while for phytogenic blends in the form of gingerol (442793), tyramine (5610), rutin (5280805), 14-deoxy-11,12-didehydroandrographiside (44575271), andrographidine E (13963769), cianidanol (9064), cyranoside (5280637), epigoitrin (3032313) and curcumin (969516). Meanwhile, data for protein modeling was obtained from Uniprot https://www.uniprot.org/ with the code Q90703 (Nitric Oxide Synthase) with the number of amino acids as much as 1136 aa (Yuan et al., 2015). The structure of protein and ligands can be seen in **Table 1**.

**Table 1.**
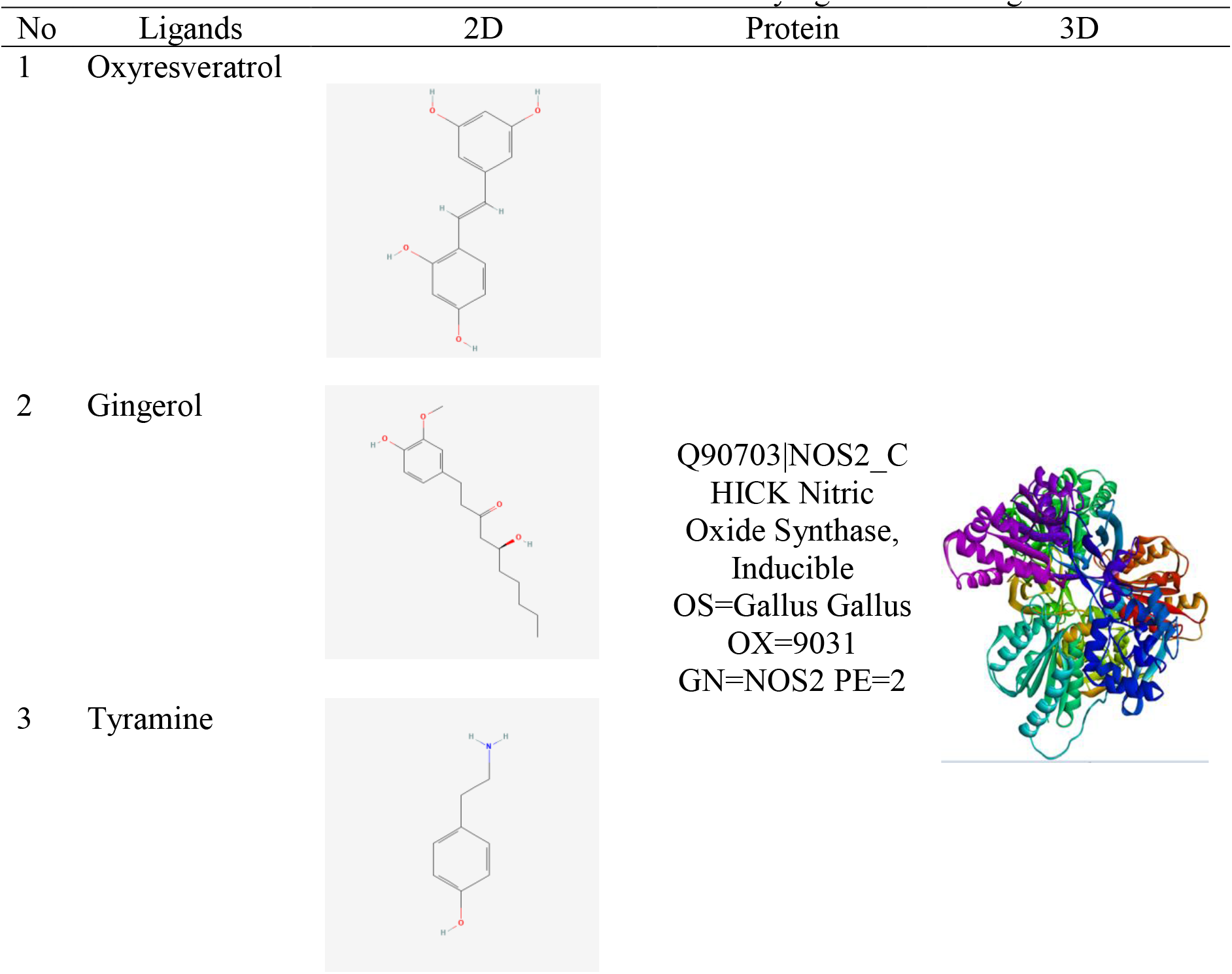
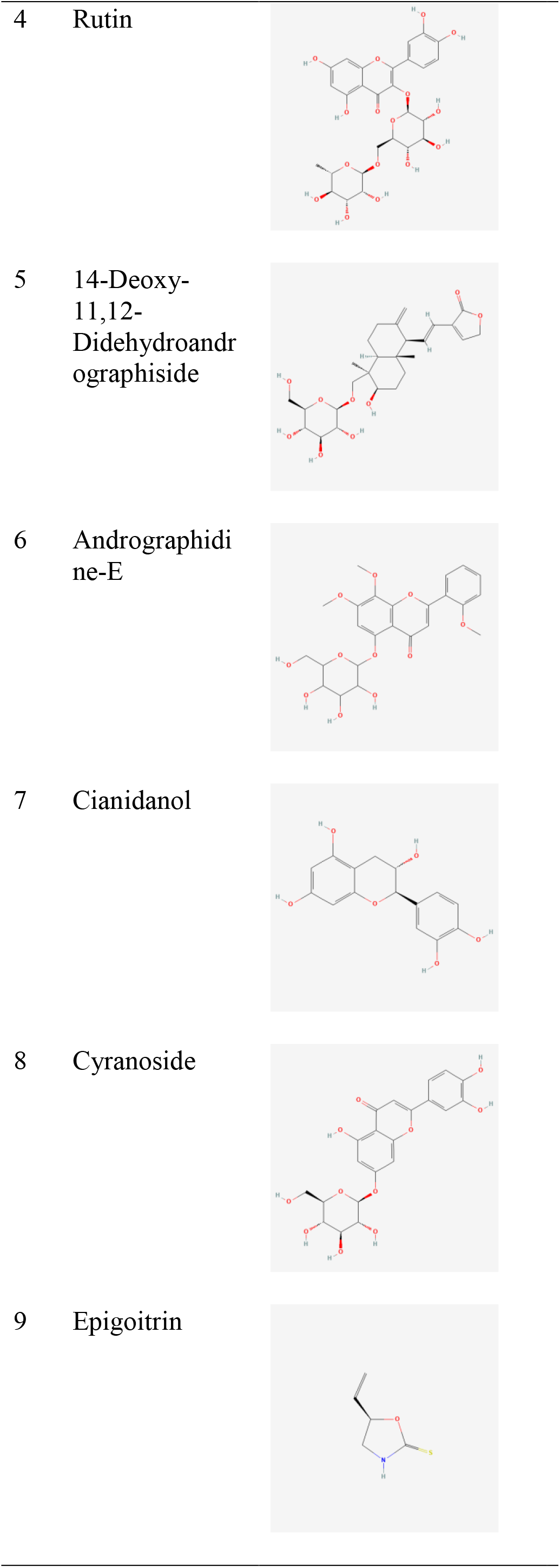
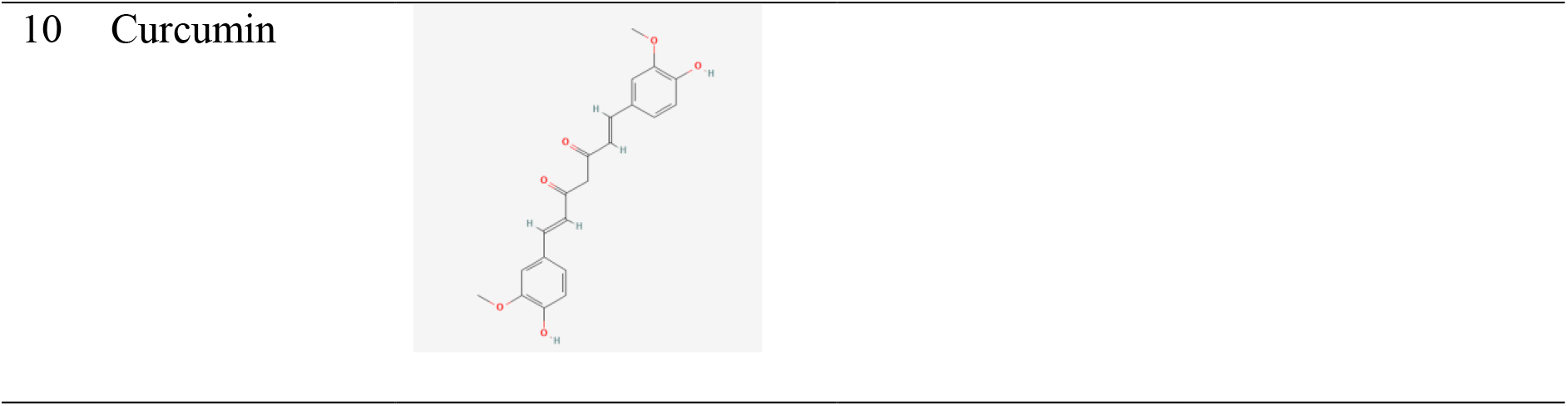
Amino acid interaction of Selected Indonesian Phytogenic Blends against iNOS

### Protein modeling and assessment

Protein modeling using SWISS-MODEL https://swissmodel.expasy.org/ by inputting protein amino acid sequences. Model selection based on coverage, GMQE, QSGE, and identity from protein template results. After the protein is successfully modeled, the assessment process uses RAMPAGE to see the protein quality based on the Ramachandran Plot.

### Molecular Docking

Protein preparation was performed using the Discovery Studio 2016 Client. Proteins are removed from water molecules and their ligands. Then ligands are prepared in Open Babel for conversion (.sdf) to (.pdbqt). Furthermore, docking uses CB-Dock 2 (https://cadd.labshare.cn/cb-dock2/php/index.php) (Liu et al., 2022). The selected docking results are the results with the first rank. Data docking analysis and visualization using Discovery Studio 2016 Client (Jiyu et al., 2019).

## RESULTS

### Protein Modeling and Assessment

Determination of the properties and functions of proteins biochemically molecular in the form of 3D protein structures can be done easily and cheaply through the in-silico method. 3D structure modeling of proteins is divided into three methods in the form of ab initio, fold recognition and homology modeling. The best 3D structure modeling of proteins whose modeling is carried out by aligning the amino acid sequence of the target protein with other proteins that have been instrumentally known 3D structure is the Homology modeling method. Templates are referred to as proteins that are already known to be 3D structures (Komari, Hadi, and Suhartono, 2020). The SWISS-MODEL server is used as a protein structure modeling because it has a high degree of efficiency so that the NOS protein can be used for further analysis. SWISS-MODEL is used by following steps in several stages consisting of determining protein target sequences, template protein identification, model creation, and model evaluation (Komari et al, 2020). The protein template used in this study refers to 7duq.1.B which is Nitric Oxide Synthase (NOS). Template selection based on coverage value, global model quality estimation of 0.73 and identity which reaches 78.39. The following is a visualization of the protein modeling presented in **Figure 1**.

**Figure 1.**
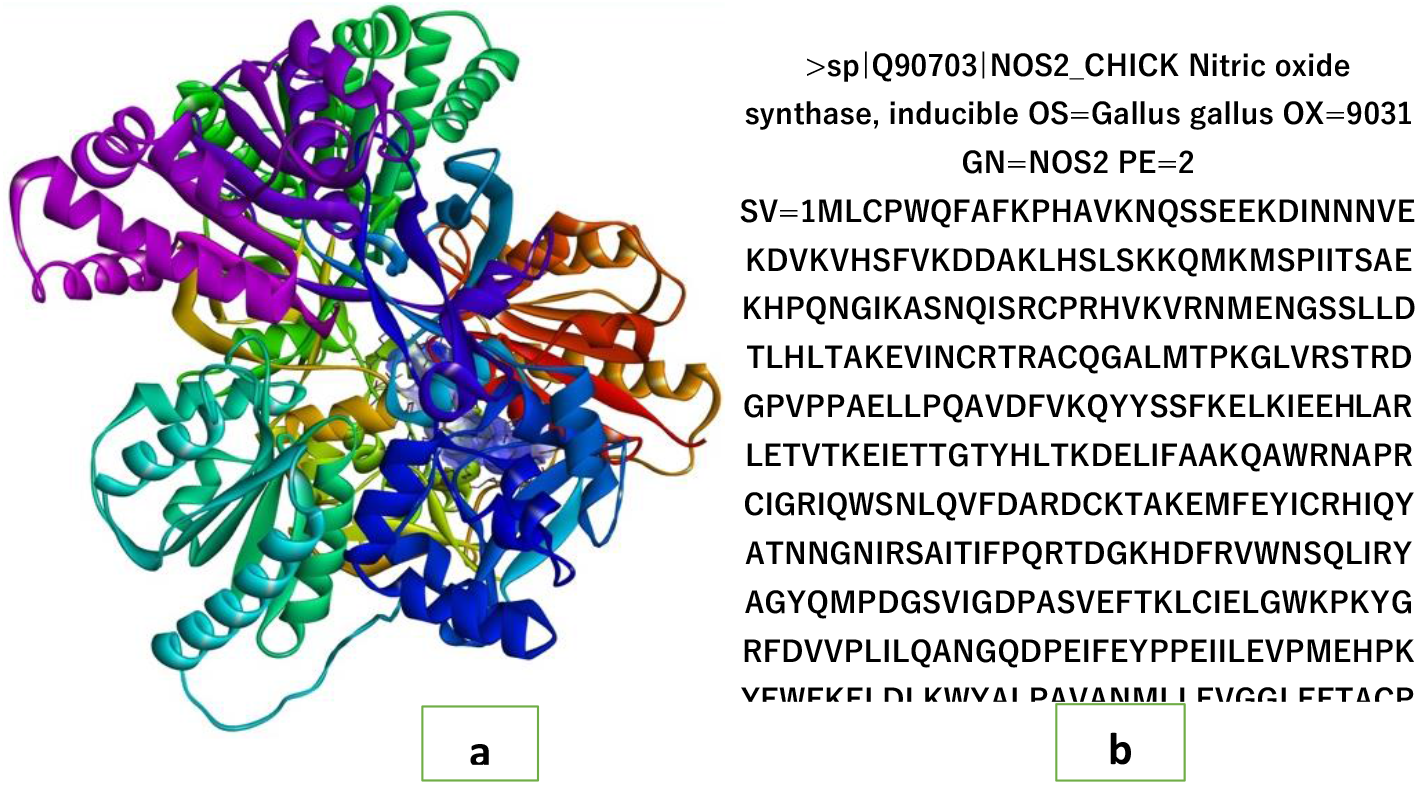
Visualization of protein modeling **Description:** (a) Protein modeling Nitric Oxide Synthase (NOS); (b) Amino acid sequences

Protein modeling data was obtained from Uniprot with the code Q90703 (Nitric oxide synthase inducible) with a total of 1136 aa amino acids found at the transcription level with the code Q90703. The protein used for modeling is a protein from poultry (Gallus gallus). The protein used is Nitric Oxide (NO) which is an important mediator of inflammation. NO is formed endogenously by the enzyme NO synthase (NOS) by utilizing L-arginine as a substrate, oxygen molecules and Nicotinamide Adenine Dinucleotide Phosphate (NADPH) as a cofactor. Most NO production is catalyzed by inducible nitric oxide synthase (iNOS). Inflammation occurs when levels of NO and iNOS levels rise in tissues (Lukiati et al., 2012).

After protein modeling, it was continued with the evaluation of the results of homology modeling using the Procheck web server which is a program for validation of protein modeling by analyzing the Ramachandran plot. The Ramachandran plot is a two-dimensional plot that describes amino acid residues in enzyme structures that have been determined through experiments into internal coordinates, where the angle Φ (phi) is the x-axis while the ψ (psi) is the y-axis divided into four quadrants. Therefore each residue can be described as one plot. Those plots that describe amino acid residues on protein structures are called Ramachandran plots. The Ramachandran plot consists of four quadrants and four regions. The four regions include most favored regions, additional allowed regions, generously allowed regions, and disallowed regions. In the Ramachandran plot, clusters formed from several residues show the secondary structure formed (Laskowski et al, 1993). The following is Ramachandran’s plot for the Nitric Oxide Synthase inducible (iNOS) protein model presented in **Figure 2**. The results of the RAMPAGE analysis show that the results of the NOS modeling protein from SWISS-MODEL are included in the category of feasible to be used in the in-silico analysis stage because highly preferred observations reached 97.368% with preferred observations of 2,350% and outliers of 0.282% as many as three amino acids, namely arginine 684, serine 697, and serine 700.

**Figure 2.**
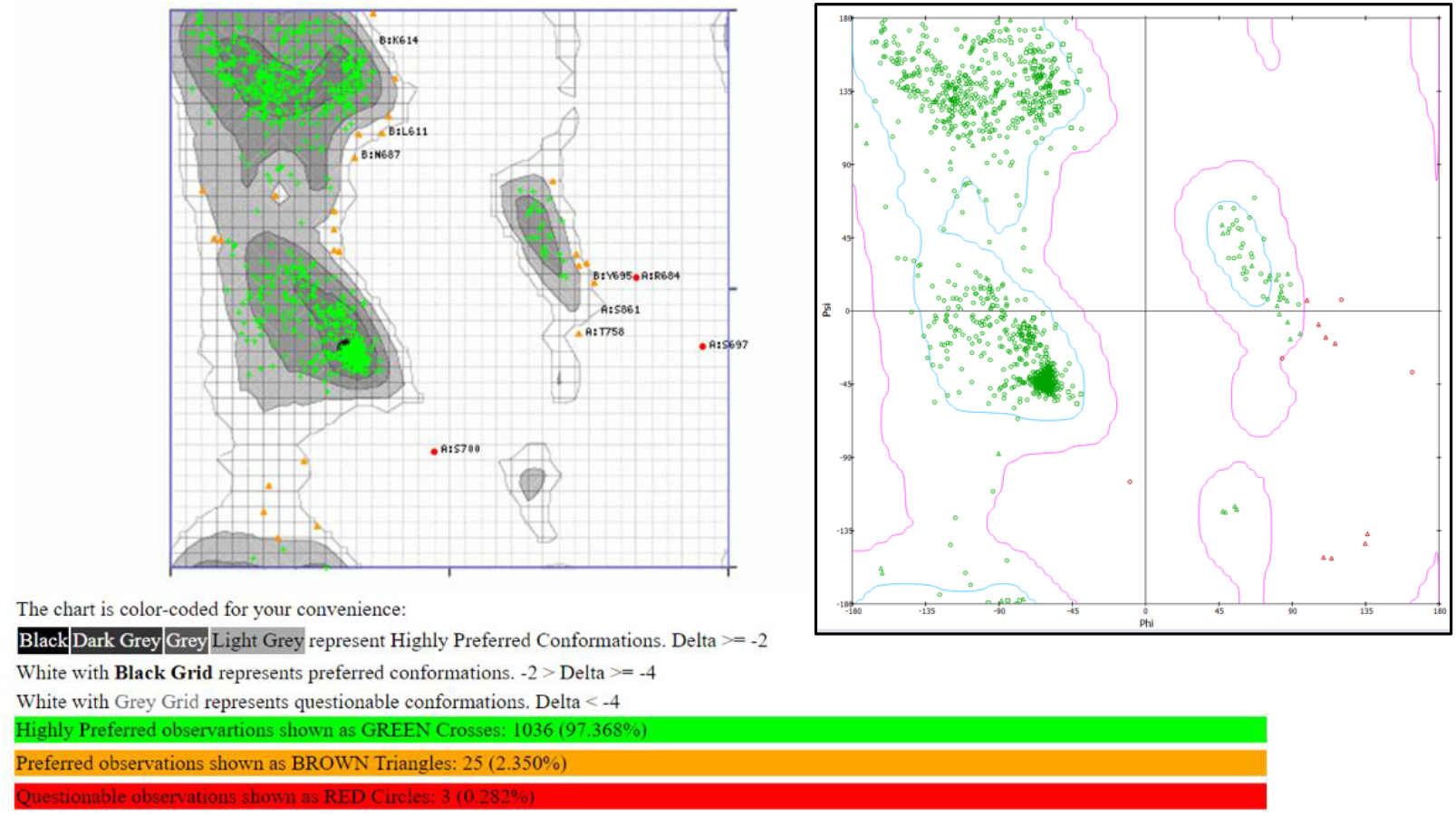
Ramachandran’s Plot for the NOS Protein Model

The main compounds that were successfully identified were continued in-silico or molecular docking tests on nitric oxide synthase (NOS) proteins computationally and validated, as listed in **Table 2**. After validation, molecular docking tests on nitric oxide synthase (NOS) proteins were performed on each major compound, Oxyresveratrol compound (control) and curcumin compound, 14-deoxy-11,12-didehydroandrographolide, andrographidine E, cianidanol/catechin, luteolin-cynaroside, tyramine, gingerol, ethyl-p methoxycinnamate/ epigoitrin (herbal waste key compound). It was found that cynaroside compounds from *Piper betle* L. with binding energy -9.4 kcal/mol, *curcuma longa* L. with binding energy -8.6 kcal/mol and *andrographis paniculata* with binding energy -9.5 kcal/mol. The three herbal plants in herbal waste can be used as a substitute for oxyresveratrol based on the value of the bond energy and the type of bond in the amino acid protein that can be seen in the figure, where the oxyresveratrol compound has a binding energy of -7.8 kcal / mol as an anti-inflammatory agent through the inhibitory pathway of nitric protein oxide synthase.

**Table 2.**
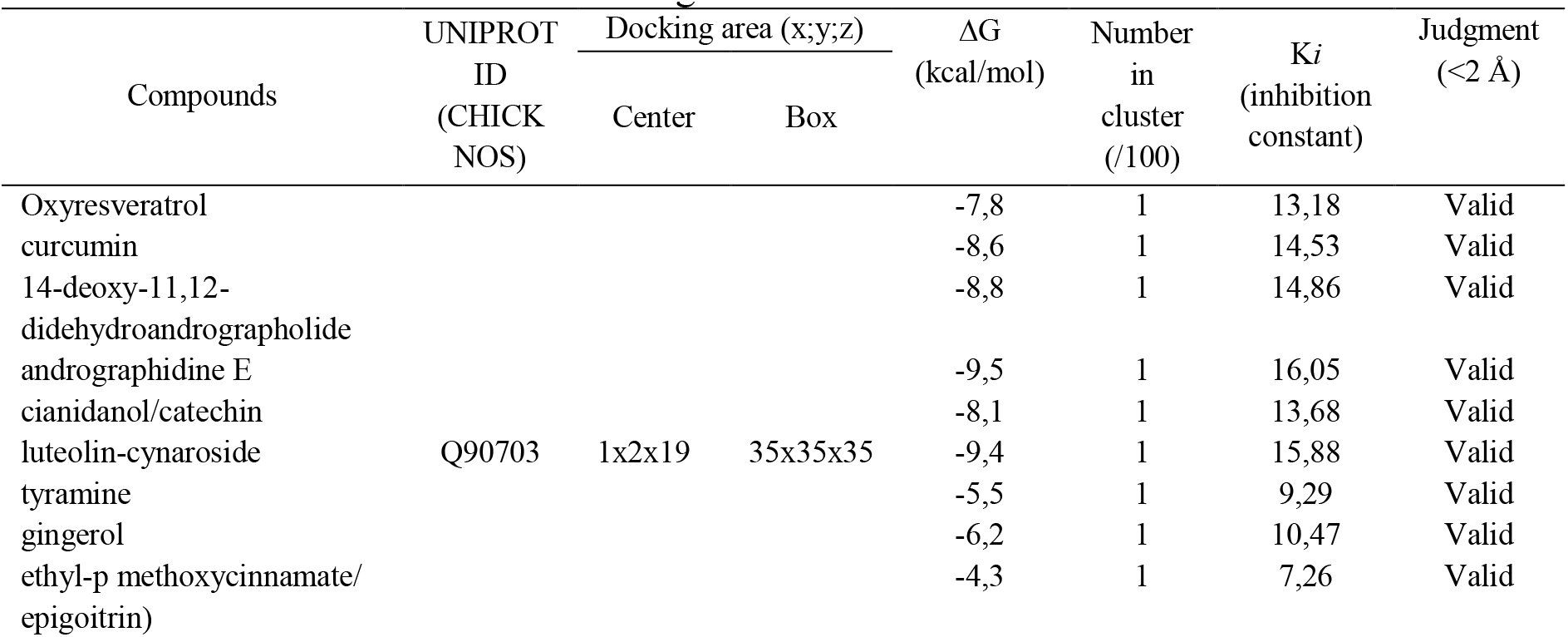
Validation of molecular docking simulation

| Compounds | UNIPROT<br>ID<br>(CHICK<br>NOS) | Docking area (x;y;z) | | $\Delta G$<br>(kcal/mol) | Number<br>in<br>cluster<br>(/100) | Ki<br>(inhibition<br>constant) | Judgment<br>($<2 \text{ \AA}$) |
| --- | --- | --- | --- | --- | --- | --- | --- |
|  |  | Center | Box |  |  |  |  |
| Oxyresveratrol | Q90703 | 1x2x19 | 35x35x35 | -7,8 | 1 | 13,18 | Valid |
| curcumin |  |  |  | -8,6 | 1 | 14,53 | Valid |
| 14-deoxy-11,12-didehydroandrographolide |  |  |  | -8,8 | 1 | 14,86 | Valid |
| andrographidine E |  |  |  | -9,5 | 1 | 16,05 | Valid |
| cianidanol/catechin |  |  |  | -8,1 | 1 | 13,68 | Valid |
| luteolin-cynaroside |  |  |  | -9,4 | 1 | 15,88 | Valid |
| tyramine |  |  |  | -5,5 | 1 | 9,29 | Valid |
| gingerol |  |  |  | -6,2 | 1 | 10,47 | Valid |
| ethyl-p methoxycinnamate/<br>epigoitrin) |  |  |  | -4,3 | 1 | 7,26 | Valid |

### Molecular *Docking*

The results of the RAMPAGE analysis show that the results of the NOS modeling protein from SWISS-MODEL are included in the category of feasible to be used in the in-silico analysis stage because highly preferred observations reached 97.368% with preferred observations of 2,350% and outliers of 0.282% as many as three amino acids, namely arginine 684, serine 697, and serine 700. The following are the results of the molecular docking from the composition of the herbal waste material used and compared with the control material.

### *Oxyresveratrol* + NOS (Control)

The hydrogen bond type control treatment has amino acid binding locations, namely GLN(A):760, GLN(B):760 and HIS(B):763. While the pi-alkyl bond is LYS(B):727, and pi-anion is GLU(A):761. While the type of pi-sigma bond is GLU(B):761 and the pi-stacked bond is PHE(B):957 (**Figure 3**). After finding the type of bond and protein in the control compound treatment, it is continued to look for compounds from herbal waste as inhibition candidates from control compounds. Indicators of the success of herbal waste compounds capable of being inhibitors in terms of the probability of binding energy values, the number of hydrogen and van der Waals bonds (Both bonds are important roles in docking results because the factors affecting protein stability.) are higher than the controls and the unfavorable bonds of donors are lower than the controls and have similarities in amino acid types with controls.

**Figure 3.**
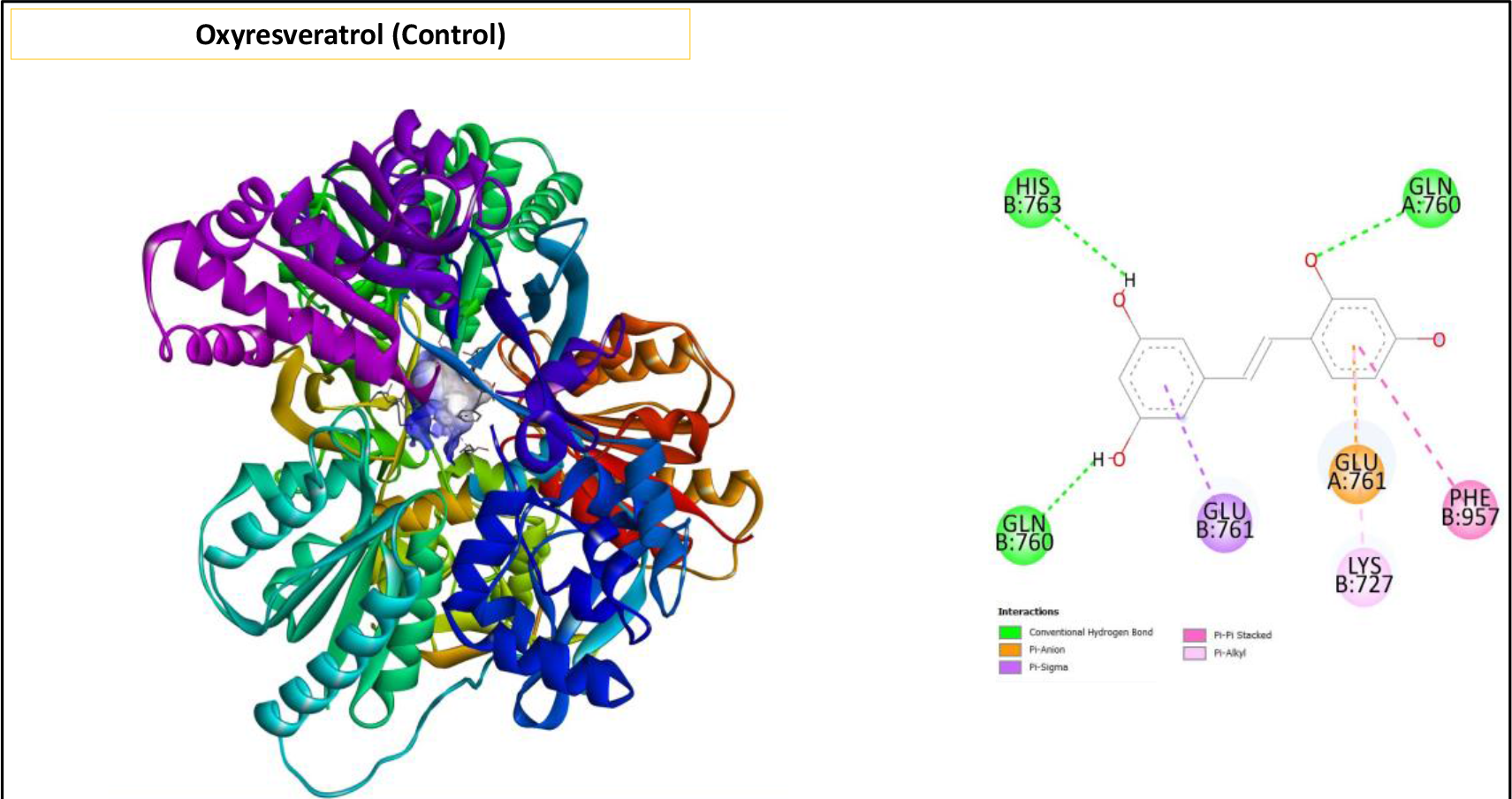
Visualization of oxyresveratrol + nitric oxide synthase (NOS) docking

Hydrogen bonding is a kind of attraction force between molecules or between dipoles that occurs between two partial electric charges with opposite polarity. Hydrogen bonds occur when molecules have N, O, and F atoms that have free electron pairs in common. The existence of hydrogen bonds, then the boiling point of H_2_O is increasing, initially the boiling point of H_2_O 100C, because of the hydrogen bond of the boiling point of H_2_O becomes 1000C. Van Der Waals bond is a bond that occurs in a noble gas that is very stable and undergoes a process so that the noble gas turns into a liquid. The larger the atomic size of a noble gas (high number of electrons) the easier it is for the gas to turn into a liquid. The donor-donor unfavorable bond (pi-alkyl) is a weak bond in protein stability. The increasing number of unfavorable bonds of donors in the docking results shows that the compound is very weak in maintaining protein stability. The following are the results of molecular docking on nine herbal waste compounds

### Nitric Oxide Synthase (NOS) Inhibition Compounds

The 14-Deoxy-11,12-didehydroandrographolide+NOS bond type is a three-bond of Conventional Hydrogen in ASN(A):597, THR(A):601, VAL(B):729, one carbon hydrogen bond in GLU(B):761 and two pi-sigma bonds in LEU(A):899, LYS(A):727 (**Figure 4a**). The bond types on the curcumin + NOS complex are four Conventional Hydrogen bonds in ASN(B):728, GLN(A):760, GLN(B):760 and SER(A):960, one hydrogen pi-donor bond in ASN(B):597 and one pi-cation bond in LYS(B):727 which can be seen at (**Figure 4b**). Conventional Hydrogen bond of cyranoside (luteolin) + NOS compounds are six Conventional Hydrogen bonds in ASN(A):728, ASN(B):728, ASP(A):570, GLU(A):761, SER(B):722 and VAL(B): 729. One Pi-Donor Hydrogen bond in ASN(A):597. One Pi-Anion Cation bond at GLU(B):761 and one Pi-Stacked bond at PHE(A):957 can be seen at (**Figure 4c**).

**Figure 4.**
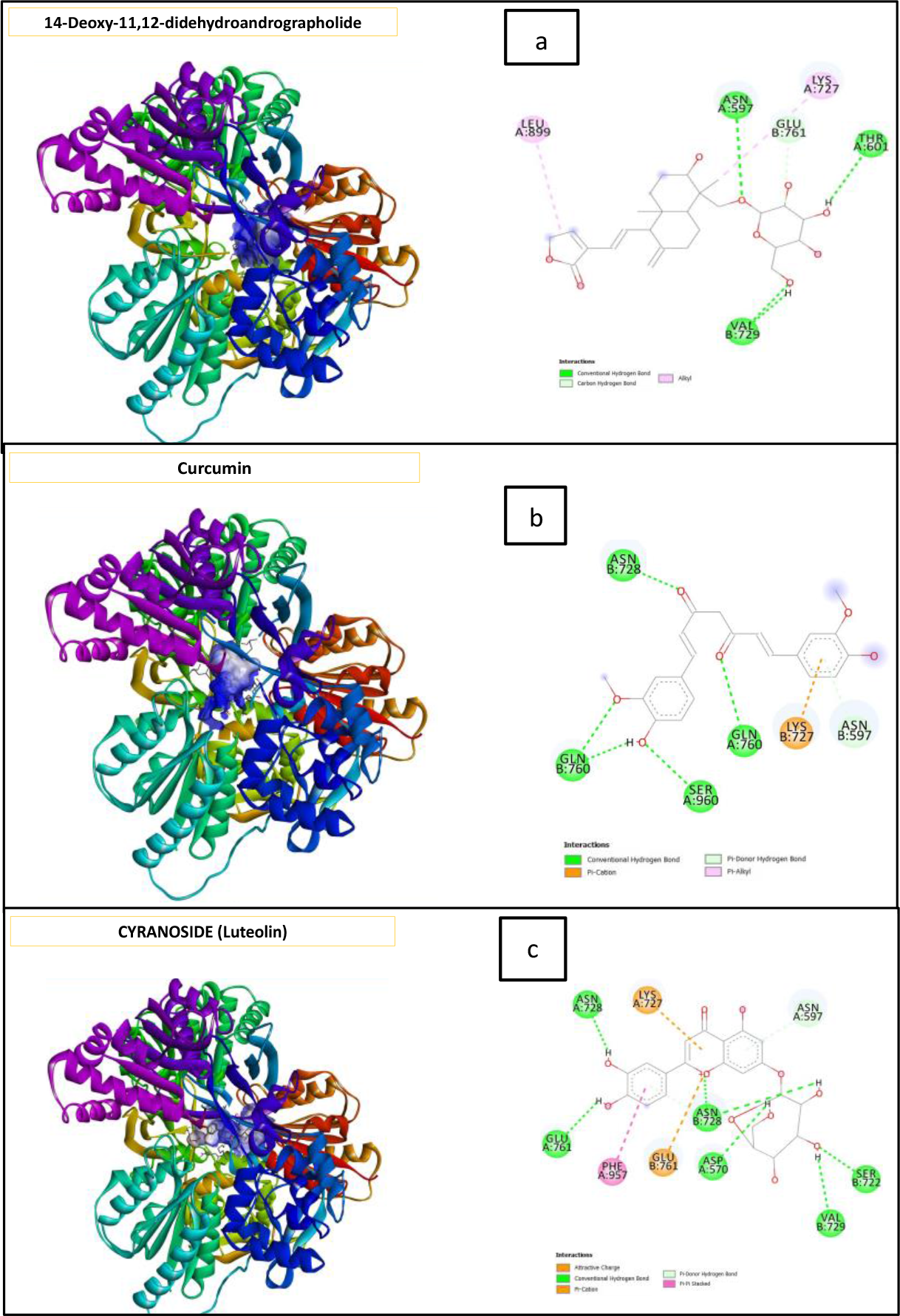
Docking Visualization **Description:** (a) 14-Deoxy-11,12-didehydroandrographolide+NOS; (b) Curcumin +NOS; (c) Cyranoside (Luteolin) + NOS

The results showed that from nine other active compounds tested, cynaroside compounds from *Piper betle* L. were found with a binding energy of -9.4 kcal / mol and a Conventional Hydrogen bond type GLU(B):761, GLU(A):761, curcumin compounds from *Curcuma longa* L. with a binding energy of -8.6 kcal/mol and a Conventional Hydrogen bond type GLN(A):760, GLN(B):760 and 14-deoxy-11,12-didehydroandrographolide compound from *Andrographis paniculata* with a binding energy of -8.8 kcal/mol and a Conventional Hydrogen glu(B):761 bond type. The three herbal plants in the herbal waste can be used as a substitute for oxyresveratrol based on the value of binding energy and the type of bond in amino acid proteins which can be seen in Figure 19, where the oxyresveratrol compound has a binding energy of -7.8 kcal / mol and the type of bond has an amino acid binding location, namely GLN (A):760, GLN(B):760 and HIS(B):763, while the pi-alkyl bond is LYS(B):727, and pi-anion, namely GLU(A):761 so that it can be used as an anti-inflammatory agent through the inhibition pathway of protein Nitric Oxide Synthase (NOS).

### Additional Compounds + Nitric Oxide Synthase (NOS)

Eight Conventional Hydrogen bonds of Rutin + NOS compounds in ASN(A):597, ASN(B):728, ASN(A):598, TYR(A):904, TYR(A):540, LYS(A):727, SER(B):722, VAL(B):729. Two Pi-Donor Hydrogen bonds in CYS(A):568, LYS(B):719, one Pi-Alkyl bond in PRO(A):902. One Alkyl bond in PRO(A):902 and one attractive change bond in ASP(A):570 can be seen at (**Figure 5a**). The andrographidine E+NOS bond type consists of several bonds, namely two carbon hydrogen bonds in TYR(A):904, ASP(B):723, four Conventional Hydrogen bonds in ASP(A):570, THR(A):601, THR(B):718, and SER(B):722, one unfavorable acceptor-acceptor in VAL(B):729, two positive-positive unfavorable bonds in LYS(A):727, ASN(A):598 and one pi-anion cation bond in GLU(A):543, can be seen at (**Figure 5b**).

**Figure 5.**
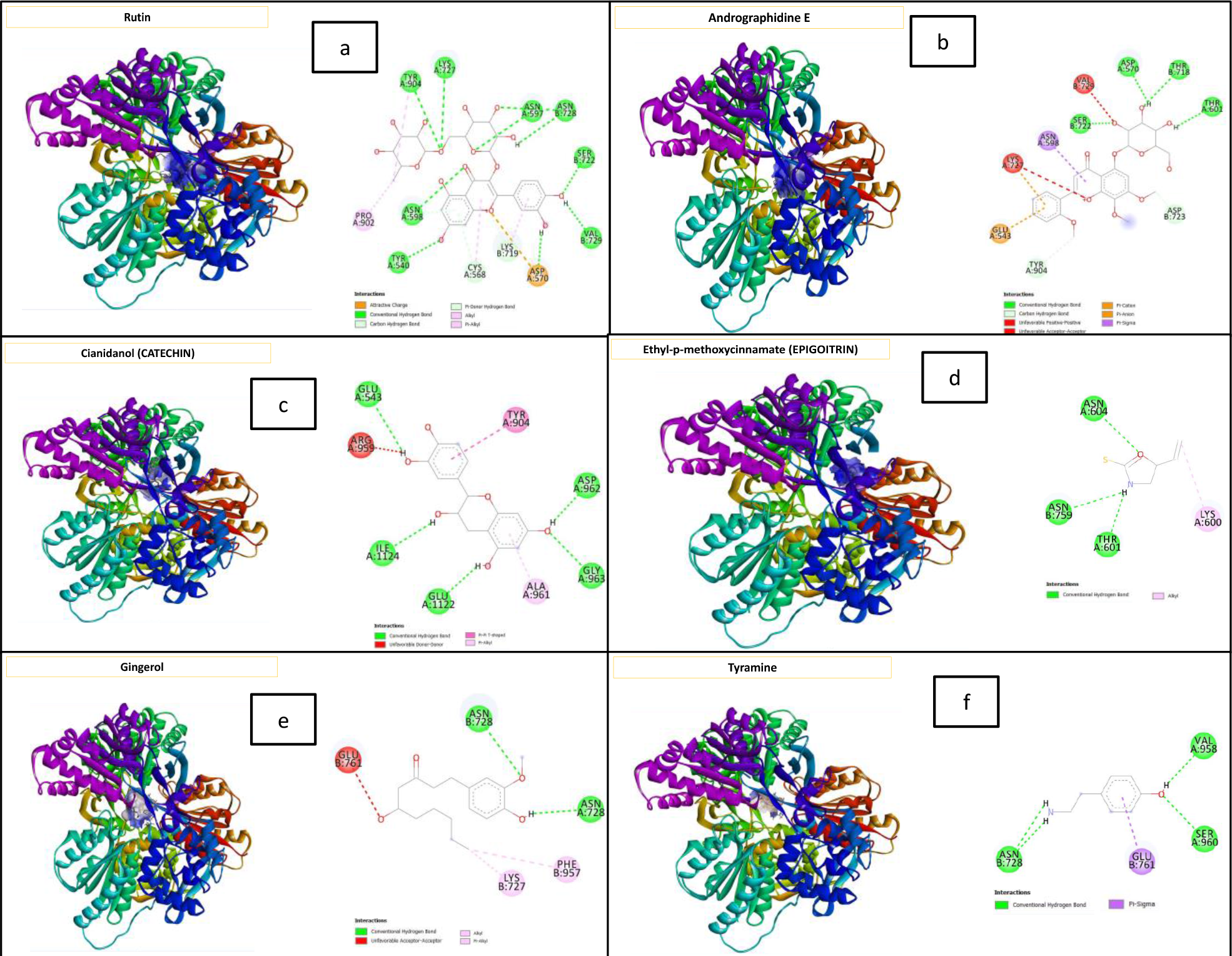
Docking Visual **Description:** (a) Rutin + NOS; (b) Andrographidine E + NOS; (c) Cianidanol (catechin) + NOS; (d) Ethyl-p-methoxycinnamate (epigoitrin) + NOS; (e) Gingerol + NOS; (f) Tyramine + NOS

**Figure 6.**
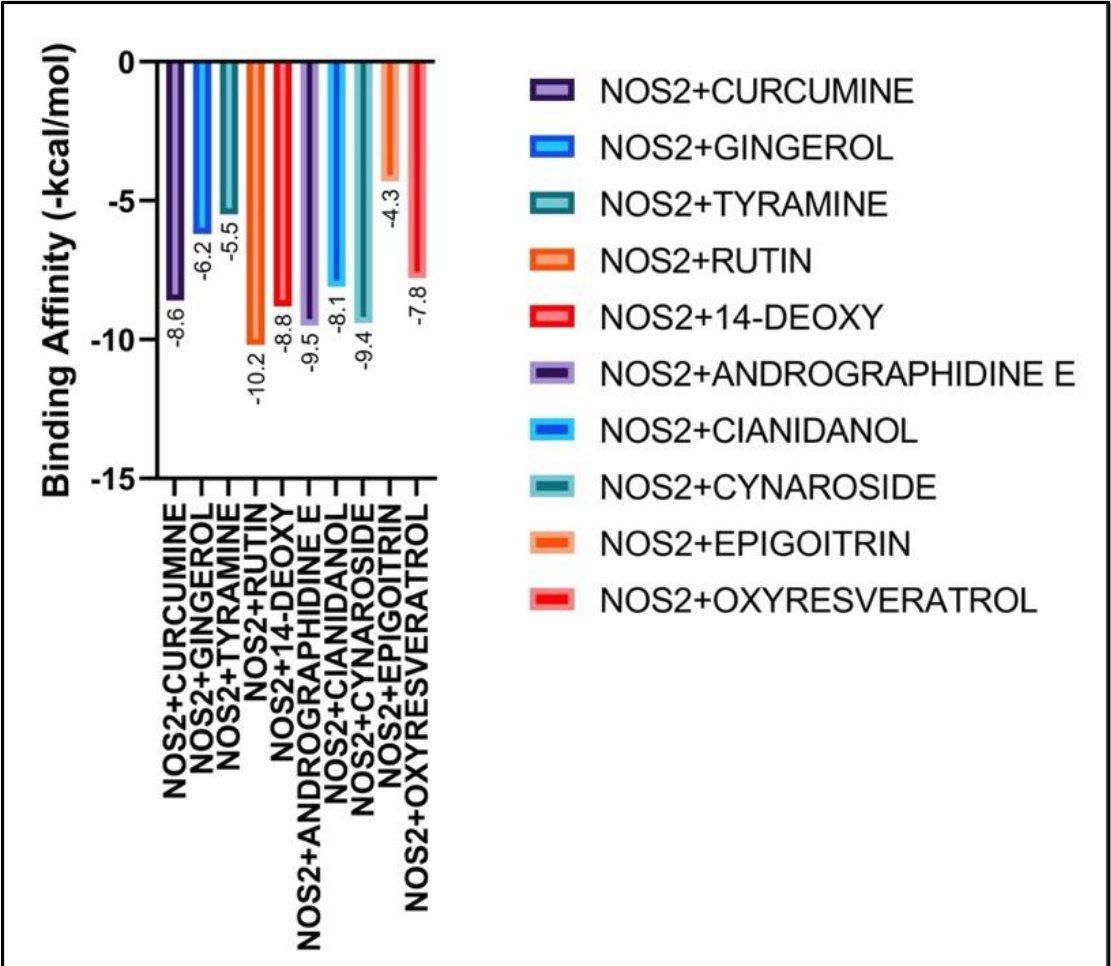
Binding energy control and phytobiotic compounds

Cianidanol (catechin) + NOS compounds have one Unfavorable Donor-Donor bond at ARG(A):959 and five Conventional Hydrogen Bond bonds at GLU(A):543, GLU(A):1122, ILE(A):1124, GLY(A):963, and ASP(A):962, one Pi-Alkyl bond at ALA(A):961 and one T-shaped Pi-Pi bond at TYR(A):904, (**Figure 5c**). The compound Ethyl-p-methoxycinnamate (epigoitrin) + NOS is located in two bonds namely one Alkyl in LYS(A):600 and three Conventional Hydrogen Bond bonds in ASN(A):604, ASN(B):759, THR(A):601 (**Figure 5d**). The two Conventional Hydrogen Bond bonds of the Gingerol + NOS compound consist of ASN(A):728, ASN(B):728), one Pi-Alkyl bond PHE(B):957, one Unfavorable Acceptor-Acceptor GLU(B):761 bond and one Alkyl bond LYS(B):727 (**Figure 5e**). Tyramine + NOS compounds have three Conventional Hydrogen bonds in ASN(B):728, SER(A):960, VAL(A):958 and one Pi-sigma bond in GLU(B):761 (**Figure 5f**).

The results showed that from nine other active compounds tested with NOS protein, it was found that five compounds from herbal waste in the form of rutin, andrographidine E, cianidanol (catechin), ethyl-p (epigoitrin), gingerol and tyramine have less probability of Conventional Hydrogen bonds and the number of Unfavorable Acceptor-Acceptor bonds is more than the control compound (oxyresveratrol) so that it has not been able to be an inhibition in suppressing inflammation in NOS protein.

### Binding Energy

From the docking results (Figure 13), data were obtained that the best bond energy possessed by rutin compounds was followed by andrographidine E, cynaroside, 14-deoxy-11,12-didehydroandrographolide, curcumin, cianidanol, oxyresveratrol (control), gingerol, tyramine and the lowest was epigoitrin.

## DISCUSSION

*Curcuma longa* L. in the form of flour can be used to optimize the work of the digestive organs to help repair body tissues and maintain the immune system of ducks and as an antioxidant agent (Rahmawati and Irawan, 2022). Apart from containing essential oils, it also contains two very important digestive enzymes, namely protease and lipase. Proteases function to break down proteins and lipases function to break down fats. Essential oils also act as bacteria destroyers and contain anti-inflammatory or anti-inflammatory properties (Rahmawati and Irawan, 2022). *Zingiber officinale* is a spice plant that is widely used because it has many benefits such as pharmacological activity, namely antibacterial, anti-inflammatory, hepatoprotector, antioxidant, immunomodulator, antihypertensive, anticancer, neuroprotector, nephroprotector, antihypertensive and anticoagulant (Adnyana and Suciyati, 2016). *Phyllanthus niruri*, L. is a medicinal plant that can be used as a phytobiotic (feed additive derived from pure plant ingredients) because it contains phytochemicals (nutrients derived from plant sources) that have antibacterial, antioxidant and effects such as alkaloids, flavonoids, saponins and tannins (Mangunwardoyo, et al., 2009) which are very effective in suppressing the growth of pathogenic bacteria. *Pipper betle* L. used as an anti-thrush, anti-inflammatory, and antiseptic. The chemical content of betel plants is saponins, flavonoids, polyphenols, and astari oil. Saponin compounds can work as antimicrobials. This compound will damage the cytoplasmic membrane and kill the cells. Flavonoid compounds are thought to have a mechanism of action of denatured bacterial cell proteins and irreparably damage cell membranes (Ajello and Susan, 2012). Essential oils from the monoterpene and sesquiterpene groups in *Kaempferia galanga* can increase endurance, appetite and antimicrobials (Male, Saputri, Qaisar and Bauang, 2012).

Homology modeling in this study is one of the tools to predict the three-dimensional structure of proteins when there is only protein sequence data on the server. Functional homology modeling to predict both structure and function. SWISS-MODEL is one of the homology modeling servers with high reliability and efficiency (Andrew et al., 2018). The results of protein modeling showed that the Nitric Oxide Synthase selected to be the model came from a model template with a PDB ID of 1tll.1.A (Nitric-oxide synthase), which had a GMQE (Global Model Quality Estimation) of 0.49 and a resolution of 2.3 Å and an identity of 51.49. GMQE is determined by a combination of the template of the crystal structure and the target sequence template that has been aligned. Next is molecular docking, where NOS protein is the target binding of control (Oxyresveratrol) and phytogenic blends from the active compounds of several plants used as feed additives in Mojosari laying ducks (*Anas javanica*) as it is known that oxyresveratrol can be used as an immunomodulator in inhibition of NOS proteins where potent blockers on NF-kB activation (Chung et al., 2003; Chen et al., 2013). The docking results showed that oxyresveratrol has a binding energy of -7.8 kcal/mol. The bond energy of oxyresveratrol in NOS proteins is strong because it consists of various bonds such as hydrogen bonds, pi-anion bonds, pi-sigma, stacked pi-pi, and pi-alkyl. This *in-silico* study aims to find an even better active compound than oxyresveratrol in NOS inhibition. The results showed that from nine other active compounds tested. It was found that the *Phyllanthus niruri* L. has a binding energy of -10.2 kcal/mol, which is the compound with the best bonding energy in nitric oxide synthase. However, compared to control, it cannot be used as a substitute for control because the interacting amino acids have nothing in common.

Similarly, andrographidine E (*Andrographis paniculata*) although it has a good bonding energy of -9.5 kcal/mol. Meanwhile, cynaroside compounds (*Piper betle* L.) can be a substitute for oxyresveratrol control with a bond energy of -9.4 kcal/mol and the similarity of amino acids that interact as many as two amino acids, namely PHE (B):363 and MET(B):229. Rutin compounds have anti-inflammatory effects, but one of the target proteins that interact with these compounds is HMGB1 (high mobility group box 1), which plays a role in acute inflammation (Yoo et al., 2014). Meanwhile, molecular docking results show that it turns out that rutin can interact with Nitric Oxide Synthase proteins as well, making this compound able to work with multi-target proteins related to inflammation. Then the andrographidine E compound also has the ability as an anti-inflammatory, where this compound can inhibit COX-2 (cyclooxygenase-2) (Jiao et al., 2019). While cynaroside compounds have 6 more hydrogen bonds than controls, research from Lee et al. (2020) supports docking results where cynaroside compounds can suppress inflammation through inhibition of iNOS, COX2, TNF-α and IL-6 protein expression in in-vitro studies. Excessive inflammation is a source of problems in poultry animals, including Mojosari laying ducks (*Anas javanica*), so the application of phytogenic blends as feed additives as anti-inflammatory agents can be further studied in vitro and in vivo stages in the future.

## Conclusion

It was found that cynaroside compounds from *Piper betle* L. with binding energy -9.4 kcal/mol, *Curcuma longa* L. with binding energy -8.6 kcal/mol and *Andrographis paniculata* with binding energy -9.5 kcal/mol. The three herbal plants in herbal waste can be used as a substitute for oxyresveratrol with a binding energy of -7.8 kcal / mol as an anti-inflammatory agent through the inhibition pathway of Nitric Oxide Synthase protein. Research related to Indonesian phytogenic mixtures as feed additives for Mojosari laying ducks (*Anas javanica*) and their ability as anti-inflammatories is being conducted in our laboratory.

## Acknowledgment

This work was supported by the Ministry of Research, Technology and Higher Education of Indonesia through the scheme of PMDSU.

## Authors Contribution

All authors contributed equally.

